# Seasonal models reveal niche changes during invasion in *Capsella bursa-pastoris*

**DOI:** 10.1101/2022.09.26.509568

**Authors:** Maya K. Wilson Brown, Emily B. Josephs

**Author notes:** Seasonal models reveal reproductive niche changes.

## Abstract

Researchers often use ecological niche models to predict where species might establish and persist under future or novel climate conditions. However, these predictive methods assume species have stable niches across time and space. Furthermore, ignoring the time of occurrence data can obscure important information about species reproduction and ultimately fitness. In this study, we generate full-year and monthly ecological niche models for *Capsella bursa-pastoris* to see if we can detect changes in the seasonal niche of the species after long-distance dispersal. We find full-year ecological niche models have low transferability across continents and there are continental differences in the climate conditions that influence the distribution of *C. bursa-pastoris*. Monthly models have greater predictive accuracy than full-year models in cooler seasons, but the inability of any model to predict summer occurrence in North America suggests a change in the seasonal niche from the native range to the non-native range. These results highlight the utility of ecological niche models at finer temporal scales in predicting species distributions and unmasking subtle patterns of evolution.

## INTRODUCTION

Species invasion of and persistence in non-native habitats affects food security, disease control, and ecosystem services (Charles and Dukes, 2007; Pyšek and Richardson, 2010; Da Re et al., 2020). Mitigating these negative effects will require predicting suitable habitat where species can invade and establish stable populations. Typically, suitable habitat is forecast using the current distribution of a species and associated measured environmental variables (Pearman et al., 2008; A. Lee-Yaw et al., 2022). Whether or not climate conditions of the native range predict the niche of the invasive range is debated, and meta-analyses have found that the realized species niches often shifts during invasion (Fitzpatrick et al., 2007; Early and Sax, 2014; Atwater et al., 2018; Liu et al., 2020; Atwater and Barney, 2021; Nguyen et al., 2022). Shifts in the realized niche could represent evolution or changes in the available environment (Bates and Bertelsmeier, 2021). However, seasonal timing can also shift between the native and non-native range, making it appear that the species niche has changed when instead only timing has changed.

Many annual plants time their lifecycle to correspond to specific seasons, so constructing niches that account for seasonal time has the potential to improve our ability to describe where and when a species will occur and detect changes in niche shifts during invasion (Hereford et al., 2017; Ponti and Sannolo, 2022). Seasonal niches are related to phenological niches, which describe where and when specific life stages occur (Grubb, 1977; Hutchinson, 1991; Wolkovich and Cleland, 2014). Determining the seasonal niche is crucial for predicting persistence, especially for annual plants who only get one chance to reproduce and cannot wait for a more favorable year (Hereford et al., 2017).

While there has been research on the phenological niches of communities of flowering plants (Bergelson and Perry, 1989; Wolkovich and Cleland, 2014; Hereford et al., 2017), we do not know how the seasonal niche can change between a native and non-native range. Comparing the seasonal niche between a species’ native range and its introduced range could reveal if and how the newly established populations are changing life-history timing in their new environments. Here, we explore the niche of *Capsella bursa-pastoris* in its native range of Europe and non-native range in North America. *C. bursa-pastoris* is a Brassiceous weed that originated in the middle east or Mediterranean region of Europe (Hurka and Neuffer, 1997). It has spread across the globe in the last 100,000 - 300,000 years (Ceplitis et al., 2005; Douglas et al., 2015), with populations in North America resulting from multiple introductions of *C. bursa-pastoris* from Europe and the Middle East in the last 600 years (Neuffer and Hurka, 1999; Wesse et al., 2021). As it colonized North America, it invaded both higher and lower latitudes than in its native range of Europe. *C. bursa-pastoris* exhibits latitudinal and elevational clines in flowering time, and some populations are more responsive than others to environmental cues for flowering (Neuffer and Hoffrogge, 1999; Cornille et al., 2022). Additionally, (Orsucci et al., 2020) showed that *C. bursa-pastoris* plants that flowered earlier than their counterparts were less affected by competition, potentially because the plants avoided interspecific competition. Most studies have explored flowering time variation in *C. bursa-pastoris* in its native range of Europe (but see (Neuffer and Hoffrogge, 1999) for flowering time variation in California). However, variation in phenological timing at a global scale is not yet known.

In this study, we investigate the niche shifts that have occurred during invasion by comparing year-long and monthly Ecological Niche Models (ENM) in the native and non-native range of *C. bursa-pastoris*. These models are commonly used to study niche differences between native and non-native populations (Liu et al., 2020; Atwater and Barney, 2021; Nguyen et al., 2022). We use the monthly models to investigate whether change in the seasonal niche is due to a shift in phenological timing or a shift in the climatic conditions that predict occurrence. If the seasonal niche of Capsella bursa-pastoris is unchanged, we might expect that the monthly model corresponding to highest abundance in Europe would have a high predictive ability on occurrences during the month of highest abundance in North America, suggesting the only difference between the seasonal niche in Europe and North America is when it occurs in the calendar year. If the abiotic factors that describe the seasonal niche are different between the native and non native range, that could suggest that plant life history has evolved over the course of invasion.

## MATERIALS AND METHODS

### Peak occurrence

For the occurrence records, we queried observations of *C. bursa-pastoris* from the Global Biodiversity Information Facility (GBIF) matching species names *Capsella bursa-pastoris (L*.*) Medik* and *Capsella bursa-pastoris L*., and where observation status was ‘present’ (GBIF.org (10 November 2020) GBIF Occurrence Download https://doi.org/10.15468/dl.nu3zhu). We removed duplicates with a matching day of observation, and coordinates (in latitude and longitude). Then, we removed nonsense observations that fell outside defined landmasses or had erroneous dates, and subsetted occurrences based on month of observation. Following Lake et al. (2020), we thinned occurrences so that no two occurred within 20 kilometers of one another to reduce the impact of spatially biased observations. In monthly models, we thinned within the month, whereas in the full year model, we thinned based on the full year dataset. We separated occurrences into a European dataset (503 - 1565 per month; mean 1002.3, sd = 136.1) and a North American dataset (32 - 288 per month; mean= 132.75, sd = 94.5).

The primary seasonal niche in each continent is defined as the month with the highest frequency of occurrences (Fig. 1). In Europe, the primary seasonal niche occurs in June; in North America it occurs in April (Supplementary Fig. 1).

**Figure 1:**
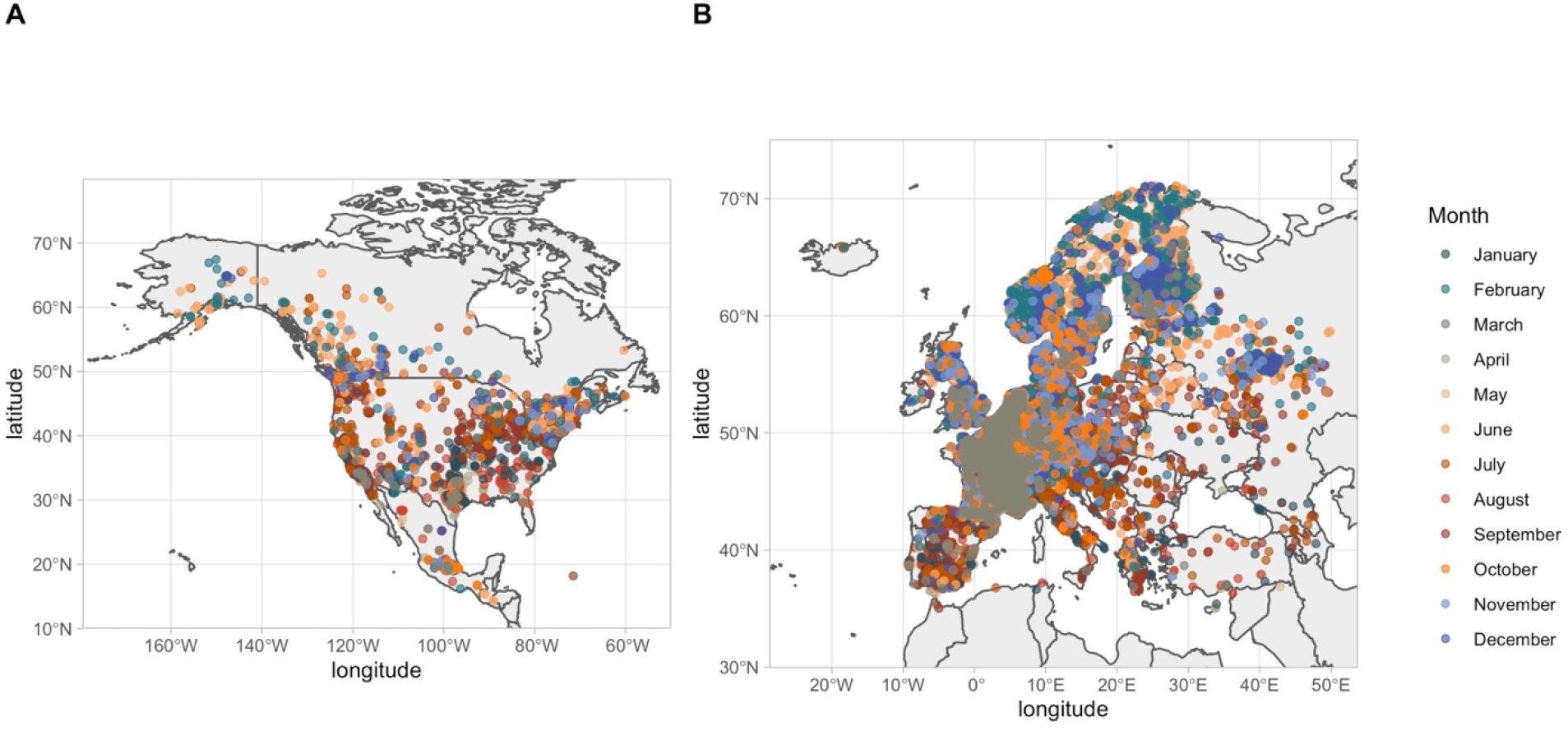
Occurrence data used in North America and Europe for this study, colored by month. Points shown are locations used to build ecological niche models after down sampling (described in methods.) Qualitatively, occurrences from the middle of the year occur at lower latitudes.

### Environmental data

We built ecological niche models for North America and Europe using data from the full year and data from each month. The full-year and monthly models use comparable environmental variables: maximum temperature of the warmest month, annual precipitation, and minimum temperature of the coldest month for full-year models and average maximum temperature, minimum temperature, precipitation, and daylength for monthly models. We obtained all environmental variables except daylength from WorldClim (representing the years 1970-2000 at a resolution of 2.5 arc-minutes). We generated an environmental predictor of daylength by calculating average daylength in each month across land masses on Earth, using latitude and the date, with the R package “geosphere’’ (Hijmans 2019). Though predictors in the full-year model and monthly models are somewhat different, they are analogous in that they coarsely capture climate in both year-long and temporal scales. Because the predictors in the full-year model represent annual averages (precipitation) and extremes (temperature), we excluded daylength as a predictor in the full-year models. Environmental features were cropped to rectangular geographic study areas around Europe and North America: 35 to 70 degrees latitude and −25 to 50 degrees longitude in Europe; 10 to 80 degrees latitude and −188 to −50 degrees longitude in North America.

### Model construction and evaluation

We used MAXENT to generate ecological niche models with default settings except that we provided background points (R package ‘dismo’; Hijmans, Phillips, Leathwick & Elith 2020). The MAXENT software contrasts the environment of occurrence records with that of pseudo-absence or background locations to build a probability distribution of species occurrences in a given study area (Phillips et al., 2017). We generated background points by picking locations proportional to the density of occurrence records for each month with the R package “KS” (Duong 2020), to account for spatial autocorrelation in occurrences and reduce the chance of an artifactually inflated omission error rate (supplementary fig. 2; supplementary fig. 3). Each model utilized a standard 10,000 background points.

For both European and North American occurrences, a geographically independent subset was withheld for model testing. We withheld occurrences within a geographic area greater than median latitude and less than median longitude of records in that month. Withheld occurrences accounted for 9-24% of all observations for each month in Europe (110-284 occurrences per month; supplementary fig. 4) and 18-44% of observations being withheld for each month in North America (9 - 68 occurrences per month; supplementary fig. 5).

We used the models to predict the niches of European *C. bursa-pastoris* in North America and evaluated their predictive ability using actual North American occurrences. We evaluated the predictive ability of all models with test observations previously withheld using a Partial Area Under the Curve (pAUC) analysis to focus on the most informative metrics of predictive ability for our study (Peterson et al., 2008), R package Kuenm: Cobos et. al. 2019). Comparing this partial AUC to a null distribution, a pAUC ratio of 1 indicates models have predictive ability equal to random; a pAUC ratio of 2 indicates models have perfect predictive ability. We chose an admissible omission error rate of 0.15 for each model and 300 bootstrapped iterations. Within each continent, we evaluated each model’s performance by using the withheld testing points for each month (ex: we evaluated the predictive accuracy of the Northern American June model on North American June occurrences withheld for testing). All models were also evaluated using the area under the receiver operator curve (AUC 0.69-0.99 in Europe, AUC 0.45 - 0.96 in North America; supplementary fig. 6).

With the European models projected to North America, we evaluated the models’ predictive ability on North American occurrences of *C. bursa-pastoris* to test for a reproductive niche shift. We used the same method described above but replaced European testing points with occurrences of *C. bursa-pastoris* in North America. Each month of North American occurrences was evaluated against all twelve models resulting in 144 evaluations of European model predictive ability on North American occurrence records.

### Climate variable influence and climate classes

To investigate how environmental features contribute to models, we extracted environmental feature contributions from each model. Additionally, we explored the direction of influence of the environmental features for each model by looking at changes in the probability of occurrences as the value of each environmental feature varied, supplied by MAXENT output. Further, we used the Koppen-Geiger climate classification (Peel et al., 2007) to better interpret climate attributes for each set of observations. Specifically, we identified the Koppen-Geiger classification for each observed plant’s location using ‘extract’ from the R package Raster (Hijmans 2021).

## RESULTS

### Models with full year of data show continental differences in environmental contributions

We generated ENMs with the full year of occurrences in Europe and North America and evaluated their predictive ability on occurrences within each continent. The predictive ability of the full year models on their own occurrences is moderate (partial AUC ratio in Europe = 1.28; partial AUC ratio = 1.32 in North America) (Fig. 2A). The European model predicts North American occurrences slightly better than it predicts European occurrences (partial AUC ratio = 1.30), whereas the North American model does not predict European occurrences better than it predicts North American occurrences (partial AUC ratio = 1.17). These results suggest that the full-year niche of *C. bursa-pastoris* has not changed between Europe and North America, but that the niche model was not very good at predicting occurrence.

**Figure 2:**
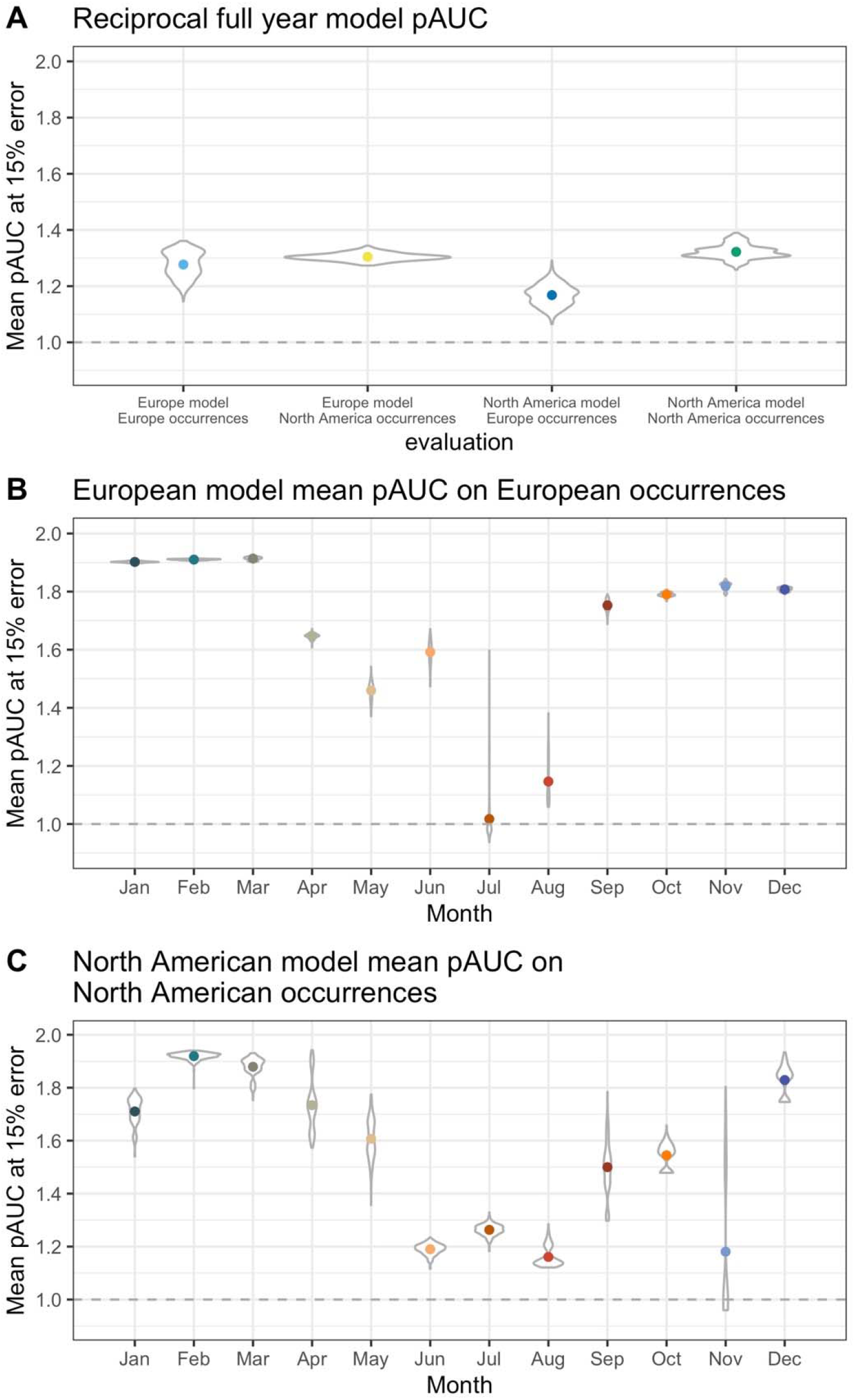
Partial area under the curve (pAUC) for each model. Means (colored points) and distribution (dark grey) from three hundred bootstrap iterations of the pAUC algorithm. A pAUC ratio of 1 indicates models have predictive ability equal to random; a pAUC ratio of 2 indicates models have perfect predictive ability.

Minimum temperature of the coldest month contributes most to the probability of occurrence of *C. bursa-pastoris* in Europe (estimated 79.2%) but minimum temperature contributes 54.7% to the model in North America. Annual precipitation contributes 4% and 33.9% in Europe and North America respectively. Maximum temperature of the warmest month contributes 16.8% to the European model and 11.3% to the North American model. Overall, these results suggest that winter temperatures may be more important for *C. bursa-pastoris* occurrence in Europe than in North America, while precipitation is more important in North America.

### Monthly models have better predictive power than year-long models

If the climate factors that affect *C. bursa-pastoris* occurrence vary by season, then models that use year-long data may fail to accurately predict occurrence. With this in mind, we generated models for occurrence by month. Additionally, we performed the analysis on a dataset where each month had an equal number of occurrences and the prediction patterns remained indicating that the results of our analyses were largely not impacted by sampling effort or data abundance (Supplementary figs. 10, 11).

Most monthly models performed well (Fig. 2) based on a Partial Area Under the Curve analysis. However, the July model in Europe is essentially equal to the null distribution (Fig. 2B) with a mean partial AUC Ratio of 1.02. The Europe August model also performed quite poorly (pAUC ratio = 1.15), potentially because of overfitting. To account for low occurrence in the testing region, we developed additional models where all months had 25% of the data withheld for testing and evaluated whether there was an effect of geography by choosing a different area to be withheld each time (Supplementary fig. 12). Increasing the amount of data withheld did not improve model accuracy for July and August substantially in any of the four scenarios, and other monthly models lost predictive ability (Supplementary fig. 12) Moreover, the September European model performed well despite having only 12% of occurrences withheld for testing (Fig. 2B), which suggests there is not a direct relationship between the proportion of occurrences withheld for testing and predictive ability in our models.

The best model performance is February and March which both have a partial AUC Ratio of 1.91. Qualitatively, model predictive ability is highest in late fall, winter, and early spring months, from September through March (Fig. 2B). In North America, predictive accuracy is best with the February model, which has a mean AUC ratio of 1.91. The August model in North America has the worst predictive ability with a mean AUC ratio of 1.16 (Fig. 2C). Generally, similar patterns hold between the seasonal predictive ability in Europe and North America, though the mid-year depression in predictive ability occurs slightly later in the calendar year in North America.

### Environmental contributions to monthly models differ between Europe and North America

Determining the environmental factors contributing to the ENM can provide information about what environments matter most for *C. bursa-pastoris* occurrence in the native and invasive ranges. In Europe, the minimum temperature of the month had the highest contribution in most months except January, July, and August. In the August model, minimum temperature and maximum temperature contribute 39.3% and 41.3%, respectively. In January daylength contributes 29.4% to the model (Fig. 3A) and minimum temperature contributes 19.8%. Otherwise, increasing minimum temperatures are positively correlated with likelihood of occurrence (cumulative log-log value), throughout the year in Europe (Supplementary fig. 7). In the European models, contributions from precipitation peak in May, and September. Maximum temperature contributes most to models in July, August, and January, and least in April. Daylength has its greatest contribution to models in the fall and winter (October through January.)

**Figure 3:**
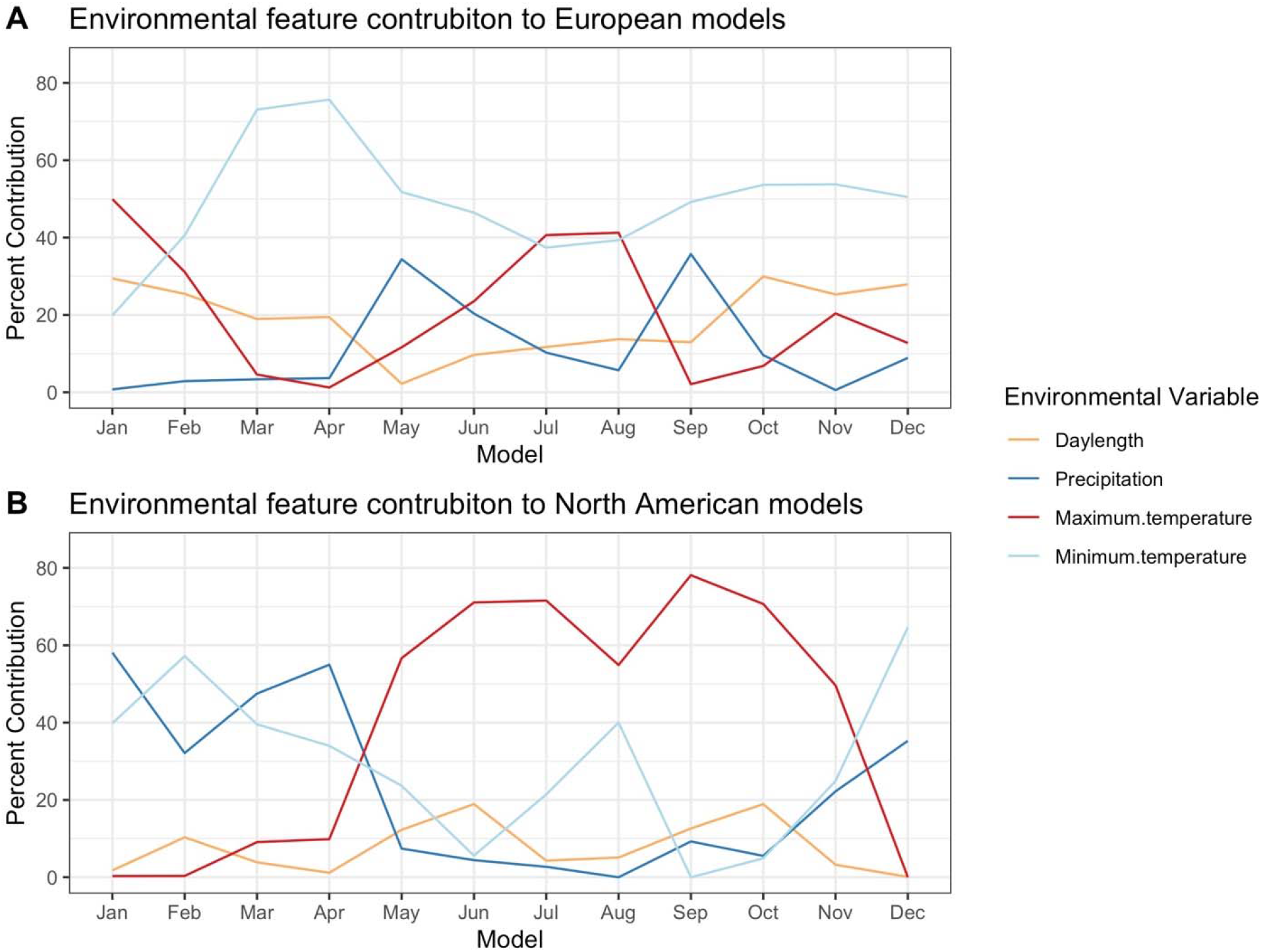
Contributions to models by environmental predictors. A) Environmental contributions to European models and B) environmental contribution to North American models throughout the year.

In North America, winter and early spring niches are most heavily influenced by minimum temperature and precipitation (Fig. 3B). However, in May, maximum temperature becomes very important and remains the largest contributor to models through November. These results suggest that cold temperatures often limit the range in Europe and that while cold temperatures are also important for winter-flowering plants in North America, hot temperatures are more important for shaping the summer range.

### European models have the highest predictive power on C. bursa-pastoris in Winter and Spring months in North America

We evaluated the predictive ability of the projections of European models on North American occurrences throughout the year to formally test if the niche changed during invasion. For each month of occurrences in Europe, we tested the predictive ability of each month’s model projected to North American climate, resulting in 144 evaluations. We interpret these projections as a probability distribution of where *C. bursa-pastoris* would occur based on the climate conditions for a particular month in Europe. For example, the August model is representative of where European August-occurring *C. bursa-pastoris* is likely to occur in North America. Shifts in the timing of life history across the range will result in shifts in the best predictive months.

Climate niches of European *C. bursa-pastoris* describe the occurrence of *C. bursa-pastoris* in North America well in some seasons. For example, the predictive ability of Fall through Winter is highest for cold months in North America (Fig. 4). Notably, *C. bursa-pastoris* that occurs in the primary seasonal niche in North America (April and May) is best predicted by the climate niches of *C. bursa-pastoris* that is abundant in Spring and Fall in Europe (Fig. 4).

**Figure 4:**
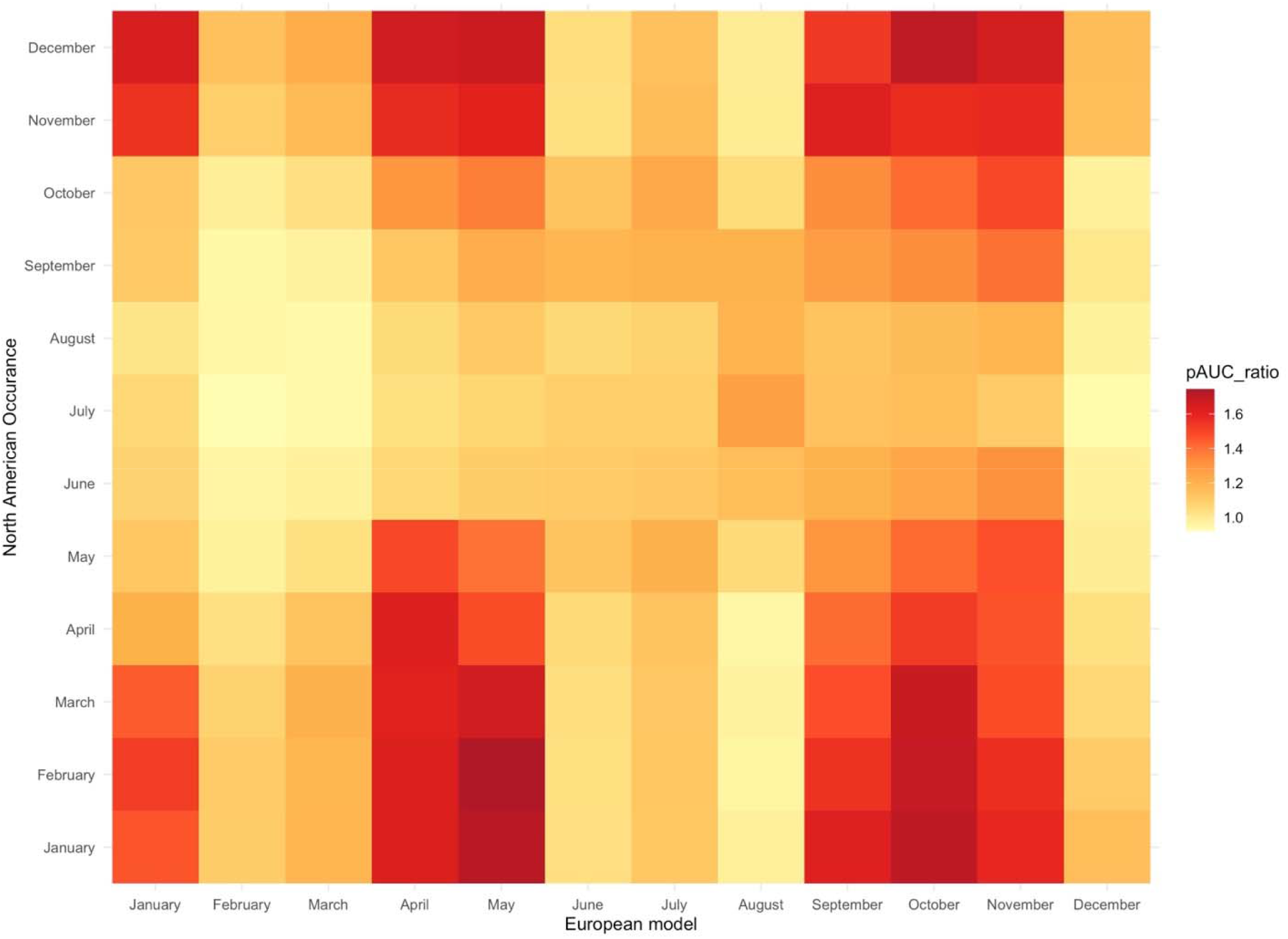
Heat map of European ecological niche model predictive ability on North American occurrences. Model of seasonal niche in Europe (June) has poor predictive ability on occurrences during peak occurrence in North America (April).

The June model, representing peak occurrence in Europe, has very poor predictive ability (AUC ratio = 1.05) on April occurrences in North America (Fig. 4). The predictive ability of European models is not dependent on the number of occurrences in predicted months (Supplementary fig. 9) Generally, *C. bursa-pastoris* that occurs in cooler months in North America (November through April) tend to be well described by European late Spring and Fall/Winter models. However, summer occurrences in North America are not well described by any model. This pattern suggests that North American summer-reproducing *C. bursa-pastoris* exist in a novel niche while North American Spring and Fall reproducing plants have a niche conserved with the native range.

### Dominant climate in each month is associated with environmental predictor contributions

We extracted Koppen-Geiger climate classes for all occurrences in the European and North American monthly models (Peel et al., 2007). In all European models the most abundant climate, given by the Koppen-Geiger climate classification system, is ‘temperate with no dry season and warm summers,’ except for the month of May, where the classification ‘cold with no dry season and warm summers’ has four more observations (Fig. 5A). Minimum temperatures contributed most to the models throughout the year, with higher minimum temperatures positively correlated with occurrences (Supplementary fig. 7). This suggests that occurrence in Europe is largely limited by freezing temperatures.

**Figure 5:**
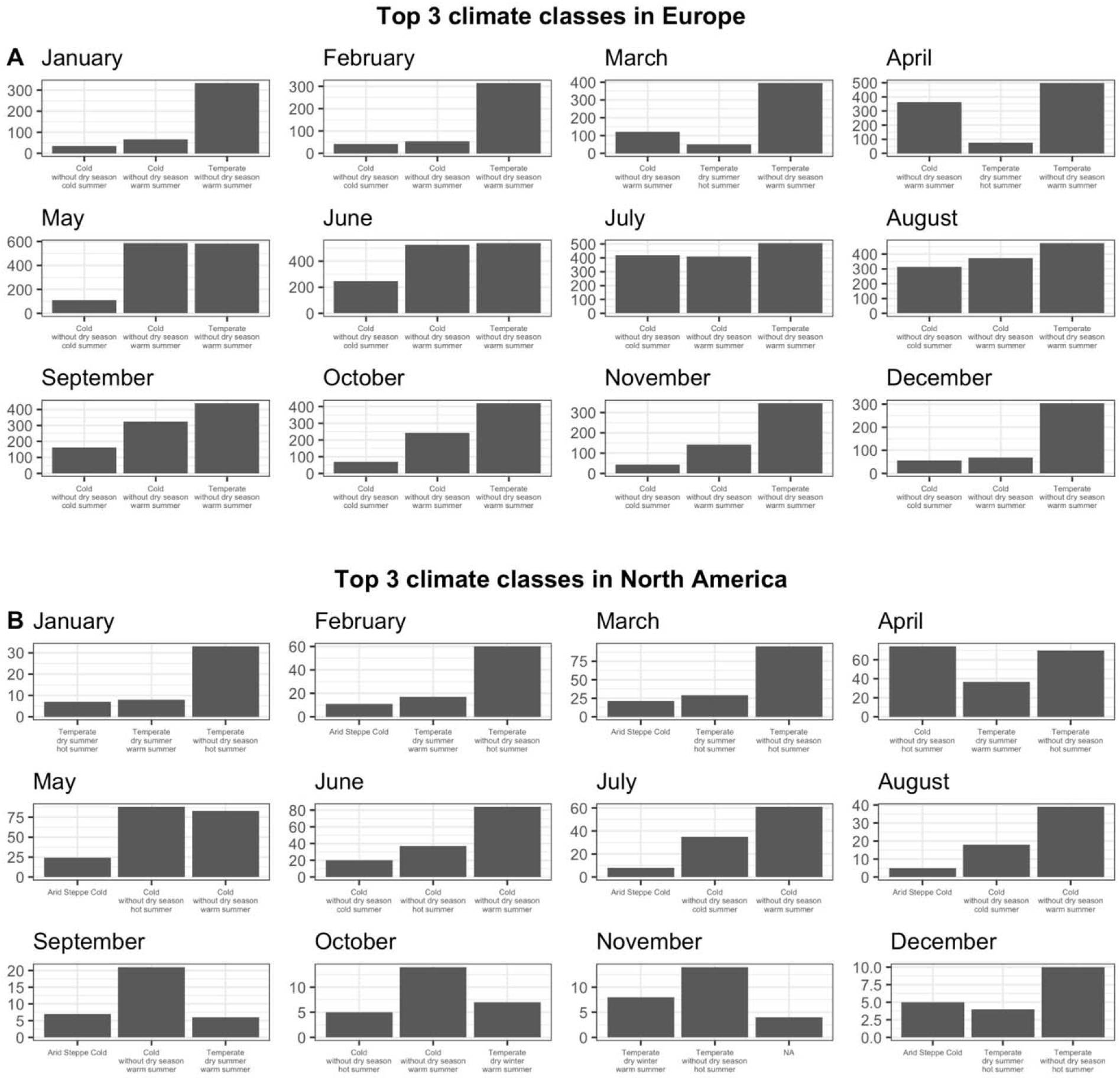
Top three climate classes for occurrences in (A) Europe and (B) North America. *C. bursa-pastoris* occurs most frequently in areas classified as ‘temperate without dry season with warm summers’ in Europe. In North America, *C. bursa-pastoris* most frequently occurs in areas classified as temperate from November to March but occurs in colder regions from April through October.

In North America, the dominant climate for *C. bursa-pastoris* is not consistent throughout the year. In Winter and early Spring (November through March), *C. bursa-pastoris* predominantly occurs in climates considered to be ‘temperate with no dry season and hot summers.’ However, from April to October, the dominant climate class is ‘cold with no dry season’ where April and May climate areas have a ‘hot summer’ and June through October locations have a ‘warm summer’ (Fig. 5B). The month of May is also when the environmental contributions to the models switch from being minimum temperature or precipitation to maximum temperature. During those months, increased maximum temperature is negatively correlated with occurrences (supplementary fig. 8). Therefore, *C. bursa-pastoris* may be limited by maximum temperature in some regions. In Europe, occurrences are always in the same climates and the abiotic factors that influence occurrence are largely the same throughout the year. But in North America, occurrences happen in different climates in different parts of the year, and those changes correspond to the changes in the abiotic factors that influence occurrence

## DISCUSSION

Here we investigated how the seasonal niche of *C. bursa-pastoris* has changed in its native European range and non-native range in North America. We found that models encompassing the primary seasonal niche in Europe are poor predictors for the seasonal niche in North America suggesting the climate niche for peak abundance has changed between Europe and North America. European models have the best predictive ability on North American occurrences in cooler months, which is unsurprising in light of the similarities between the uropean climate and North American climate from November to March. It could be that these *C. bursa-pastoris* populations have responded plastically to the same environmental predictors for seasonal changes, so they are able to match the niche from their native range. However, summer occurrence of *C. bursa-pastoris* in North America is not well-predicted by European summer models, suggesting that *C. bursa-pastoris* has moved into a new seasonal niche.

The relative contributions of environmental predictors in European and North American models during the summer highlights differences in the abiotic environment important for *C. bursa-pastoris* between the two continents. While the patterns of what environmental contributions contributed to each model are complex, there is a general trend that minimum temperature is more important in Europe while maximum temperature is more important in North America. A similar pattern is seen in Arabidopsis *thaliana*, a close relative of *C. bursa-pastoris* (Yim et. al., 2022). Our results suggest that, despite having similar climates in the native and non-native range during some parts of the year, different selective pressures could shape phenology as *C. bursa-pastoris* dispersed into other climates in North America.

One might assume the differences in timing of the seasonal niche between Europe and North America are primarily driven by differences in latitude. If this were the case, we would expect to see that a given European monthly model has high predictive ability on North American occurrences from preceding months. In effect, we would expect there to be a deeper toned diagonal line through the Figure 4, but southeast of the x=y line, indicating earlier phenology in North American because of the continent’s lower mean latitude. Given our results, it is unlikely that the differences in the seasonal niche of *Capsella bursa-pastoris* are due largely to differences in latitude.

The ability of *C. bursa-pastoris* to shift its phenology in the non-native range, either through plasticity or rapid adaptation, could be responsible for *C. bursa-pastoris’* large distribution. Our work showing shifts in the seasonal niche fit in with other work on phenology in *C. bursa-pastoris*. For example, common garden experiments in *C. bursa-pastoris* have shown that phenology is plastic but also has genetic variation across the range (Cornille et al., 2022) and that phenology shifts affect competition, potentially allowing *C. bursa-pastoris* to persist in highly-competitive environments (Orsucci et al., 2020). Understanding the mechanism through which *C. bursa-pastoris* has managed to quickly expand its range could reveal broader patterns of evolution in invasive species, especially other polyploids and selfers.

Using species occurrences as a proxy for presence has limitations, mainly that observations could be biased towards human activity; georeferenced observations of *C. bursa-pastoris* are likely to be spatiotemporally correlated with sampling effort. We attempt to account for spatial correlation between sampling effort and environmental predictors by thinning occurrences to be no closer than 20km to one another. There is also a considerable skew between the number of occurrences per month in Europe versus North America (mean per month before downsampling in Europe = 8087.17 occurrences; North America = 246 occurrences). We cannot completely rule out the possibility that the frequency of georeferenced observations in each month are correlated with sampling effort, but the results of models with equal numbers of occurrences per month suggest sampling effort does not largely impact our results (Supplementary figs. 10 and 11). Nonetheless, ecological niche models of the seasonal niche of *C. bursa-pastoris* in Europe are poor predictors of the seasonal niche in North America. Given the similarity between the climates of North America and Europe and fidelity in model prediction, there is evidence that the seasonal niche of *C. bursa-pastoris* evolved with long-distance dispersal. The seasonal niche of *C. bursa-pastoris* in Europe best predicts North American *C. bursa-pastoris* occurrences in cooler months. Notably, the environmental pressures important for determining the seasonal niche differ between Europe and North America, but on both continents, temperature appears to have a large influence. Experimental manipulations could further elucidate this relationship (as in Querns et al. (2022)). Additionally, it is imperative to expand populations used in experimental trials beyond those in Europe and North America to include populations where *C. bursa-pastoris* has established in increasingly disparate climates.

## CONCLUSIONS

The use of ecological niche models at fine temporal scales has the potential to uncover changes in phenology that directly impact fitness. Our results indicate that there have already been changes to the seasonal niche of *C. bursa-pastoris* after long-distance dispersal from Europe to North America. Given the broad similarities in climate between Europe and North America, it is possible that populations of *C. bursa-pastoris* persisting in very different climates with different demographic histories show even stronger signals of change. Understanding how geographically isolated populations of *C. bursa-pastoris* respond to climate that ultimately influences fitness will provide invaluable insight into how plants may respond to climate change.

## Acknowledgements

We thank Seema Sheth and Jesse Lasky for helpful comments. This work was supported by the National Science Foundation Graduate Research Fellowship under Grant No. DGE-1848739 to M.K.W.B. and NIH-1R35GM142829 to E.B.J, as well as funding from Michigan State University. Computational resources from the Institute for Cyber-Enabled Research at Michigan State University.

## Author Contributions

MWB and EBJ designed the project and wrote the manuscript. MWB did analyses.

## Data Availability

Data available on Dryad:

## Supplementary Figures

**Supplementary figure 1:**
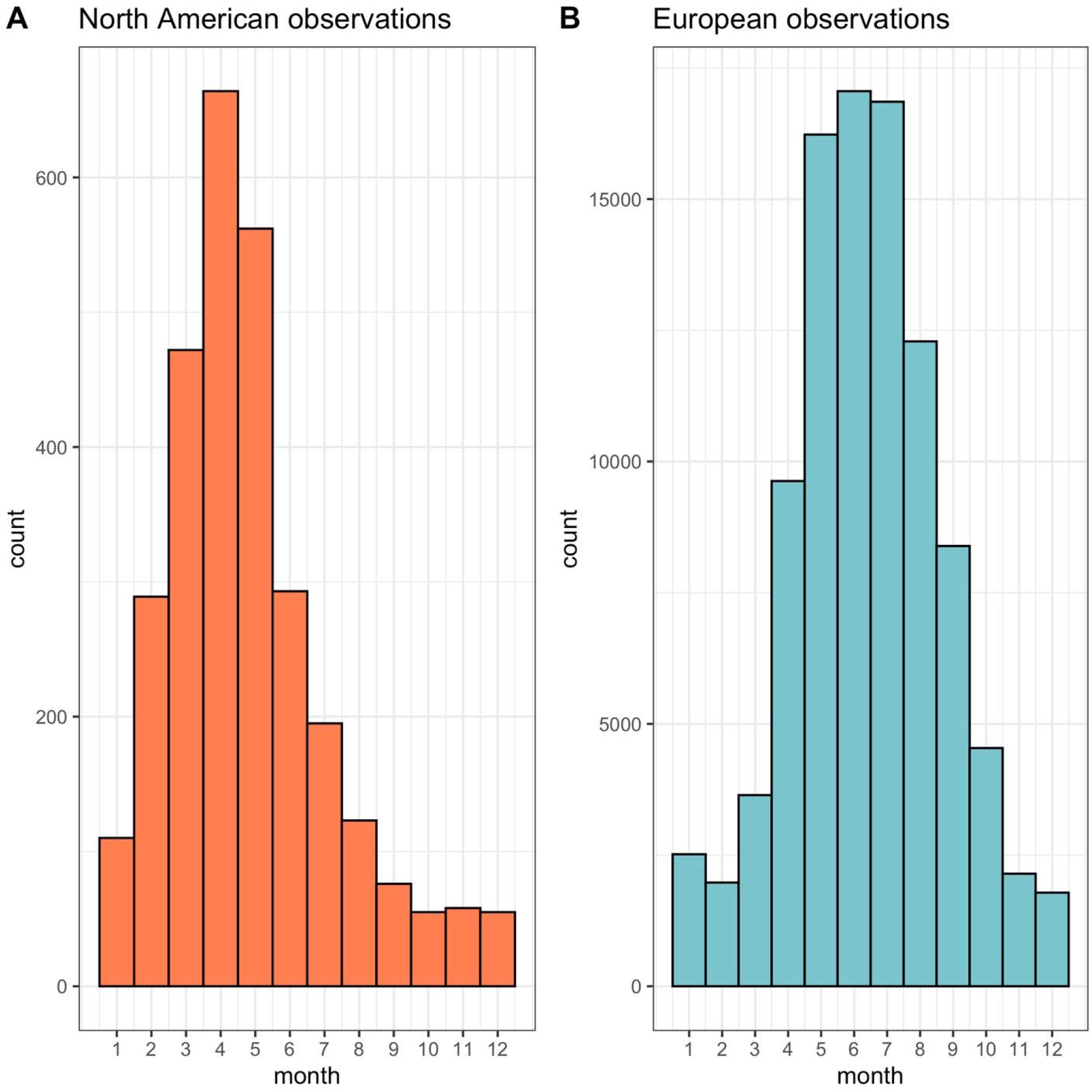
Seasonal niche in North America (A) and Europe (B) as defined by number of occurrences.

**Supplementary Figure 2:**
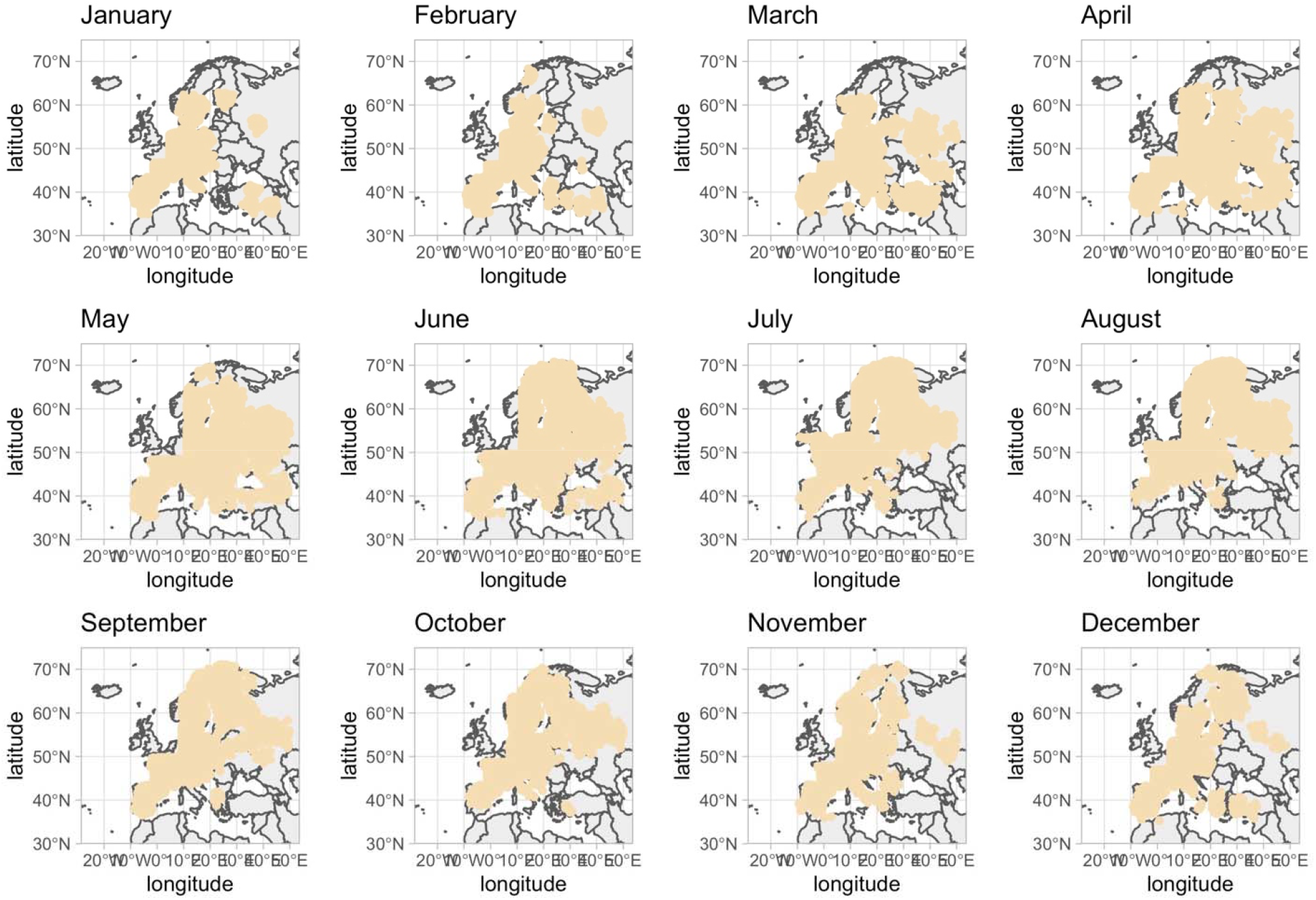
Background points used in monthly models for Europe

**Supplementary Figure 3:**
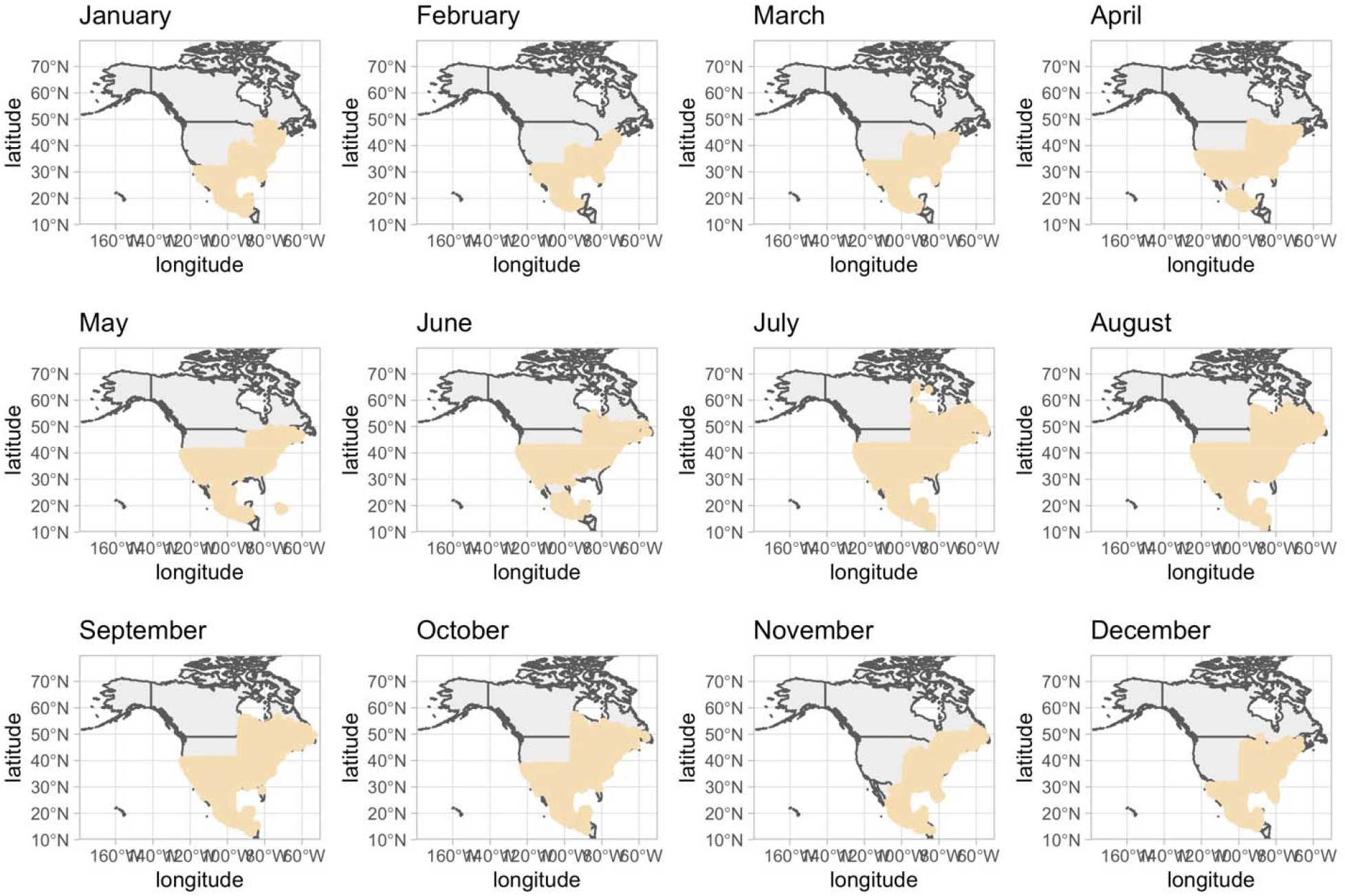
Background points used in monthly models for North America

**Supplementary Figure 4:**
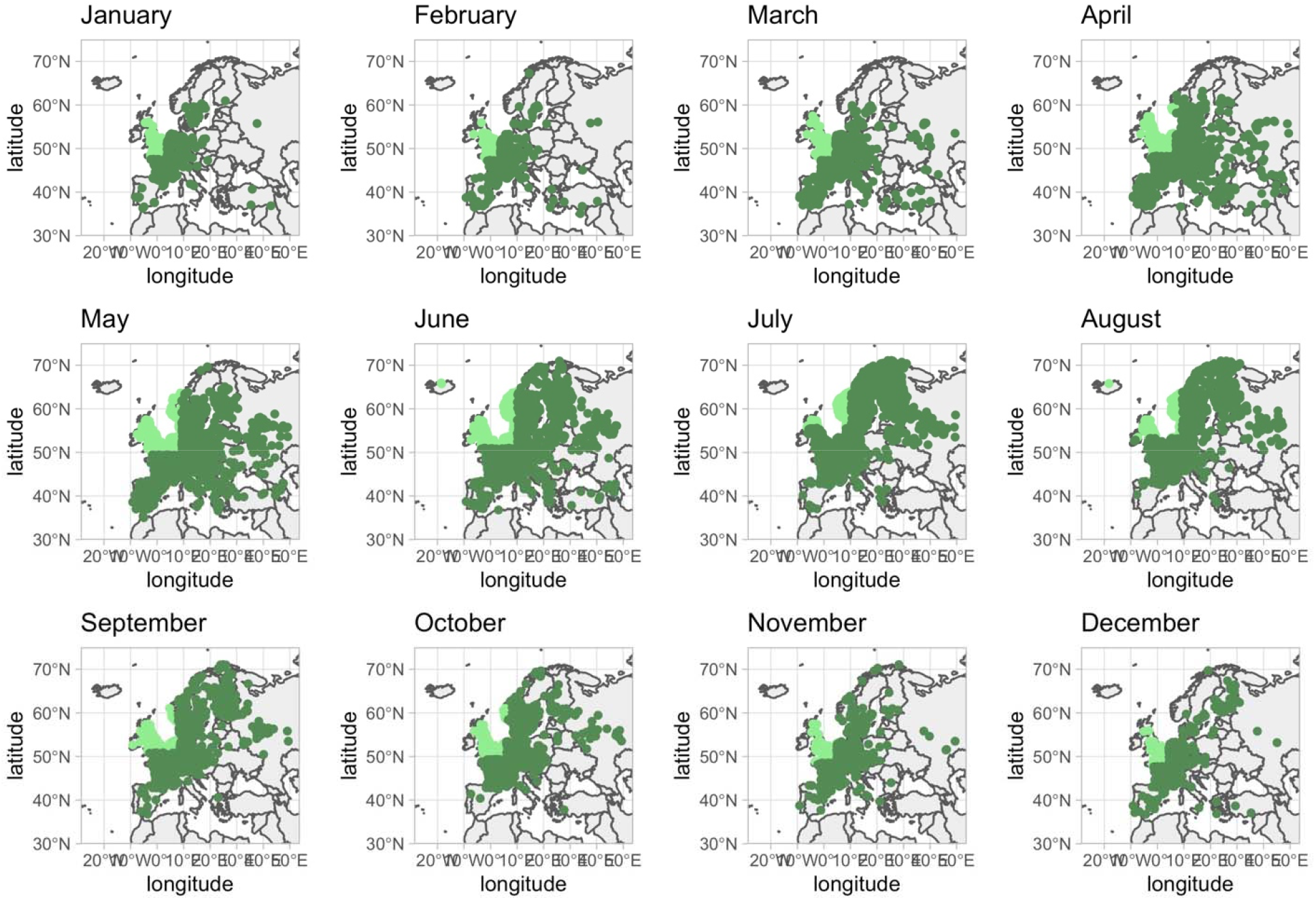
Testing (light green) and training (dark green) points in Europe

**Supplementary Figure 5:**
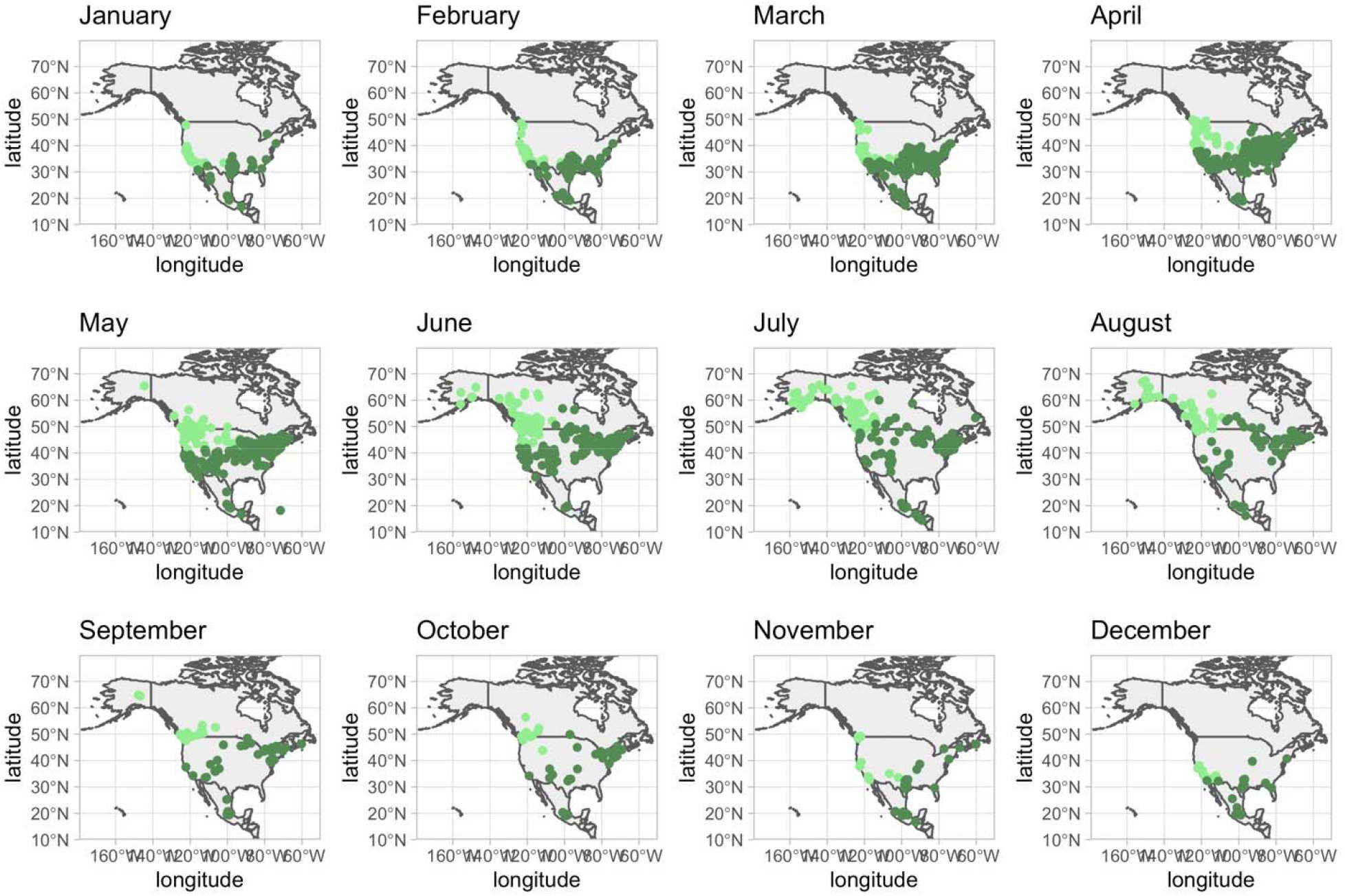
Testing (light green) and training (dark green) points in North America

**Supplementary Figure 6:**
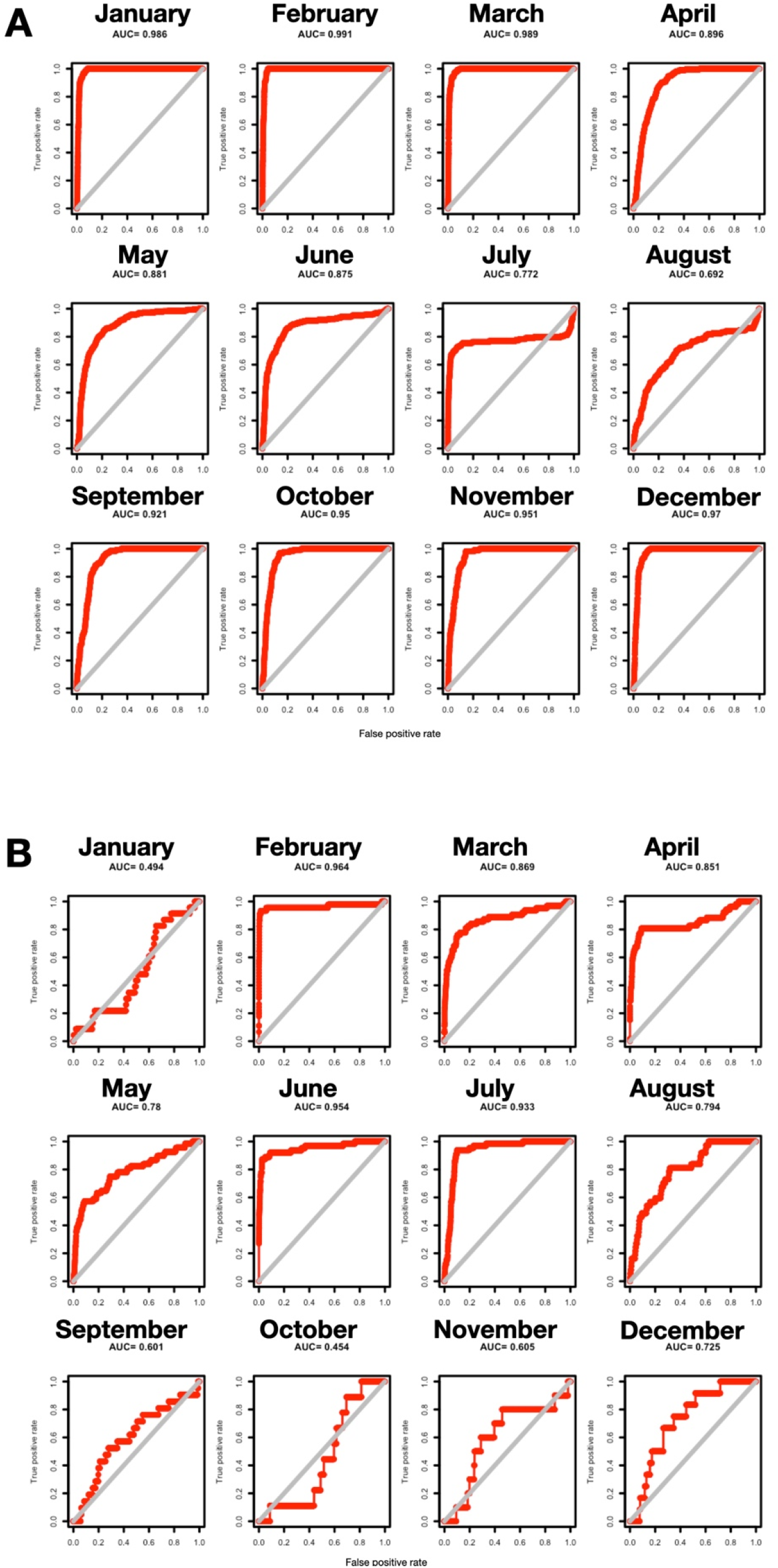
Standard ROC curves for monthly models in (A) Europe and (B) North America.

**Supplementary Figure 7:**
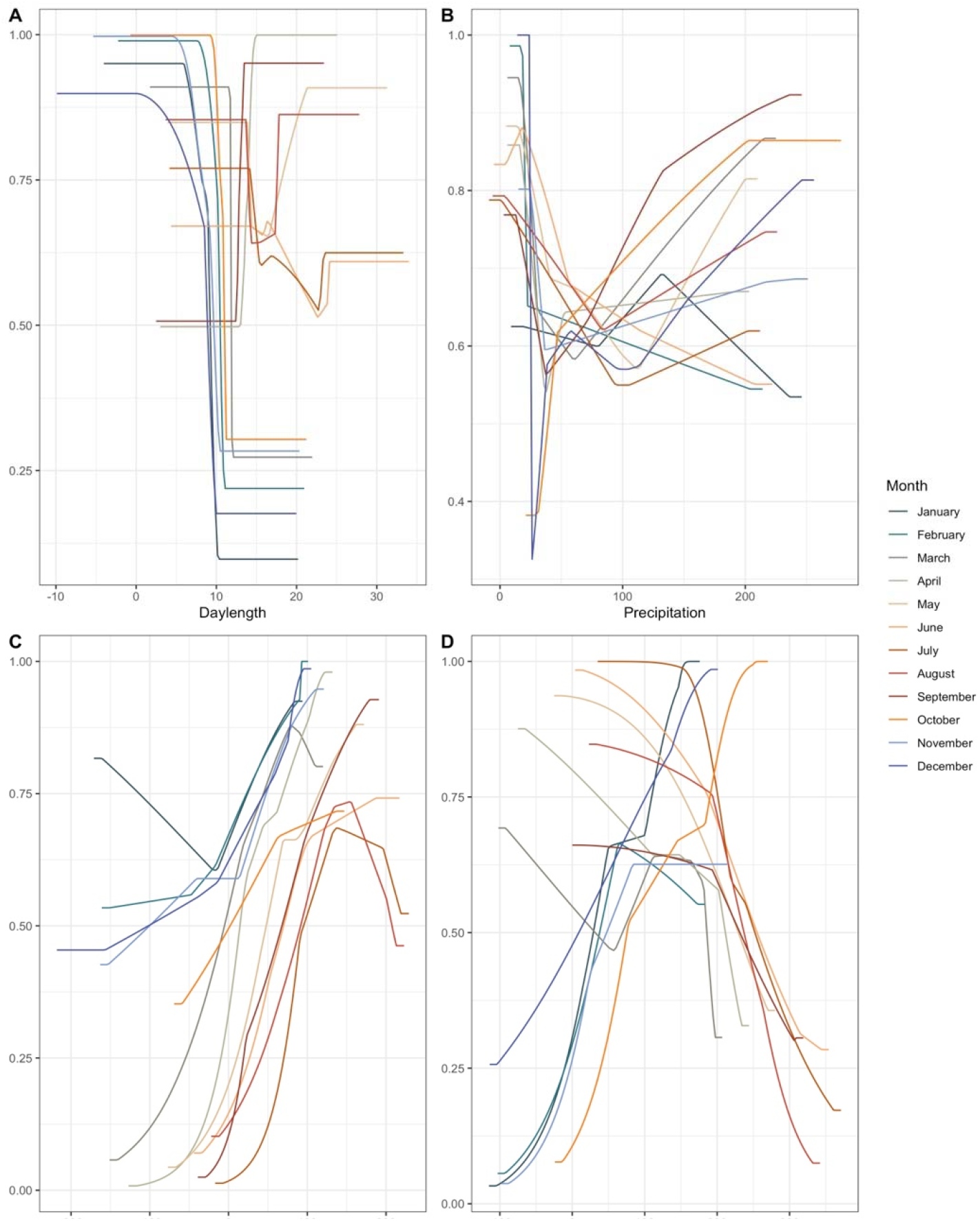
Cumulative log-log likelihood of occurrence across environmental gradients for Europe

**Supplementary Figure 8:**
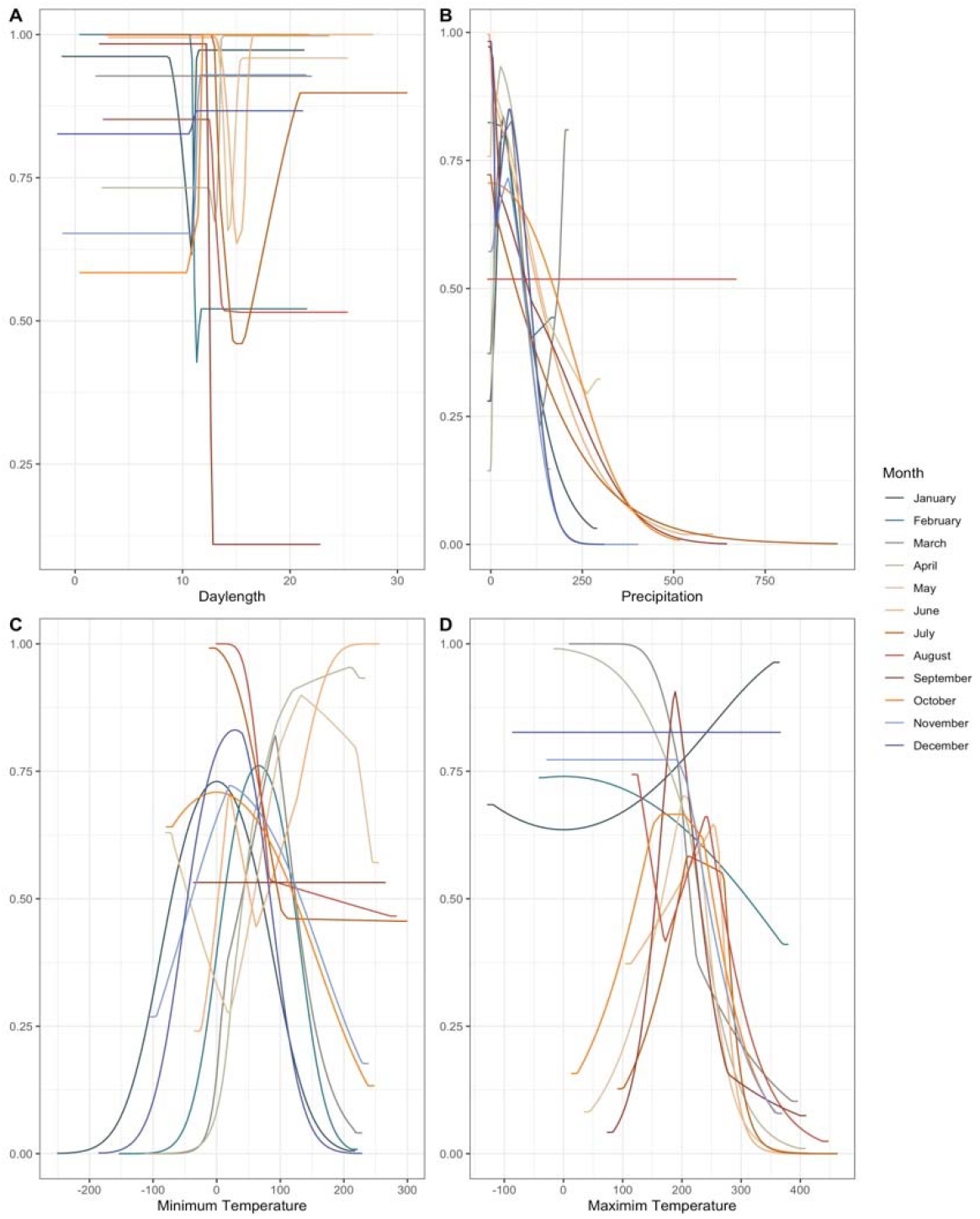
Cumulative log-log likelihood of occurrence across environmental gradients for North America

**Supplementary Figure 9:**
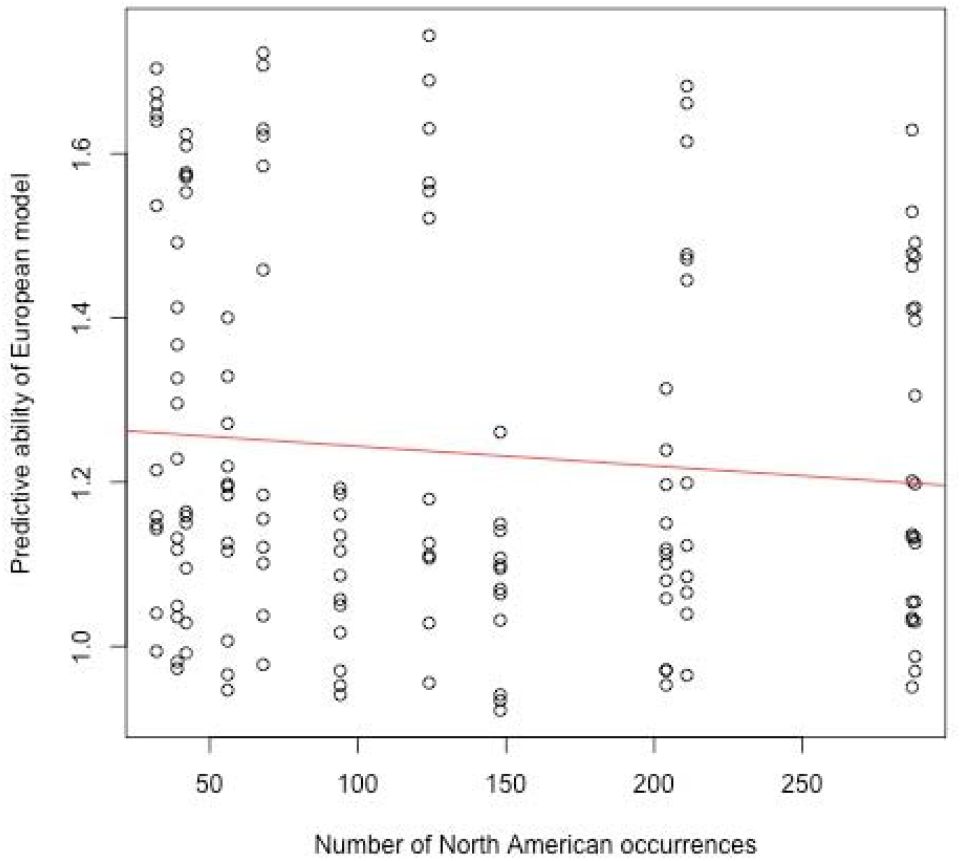
The predictive ability of European models is not significantly associated with the number of occurrences being predicted in North America. Red line is a liner model where the slope has a p-value of 0.268.

**Supplementary Figure 10:**
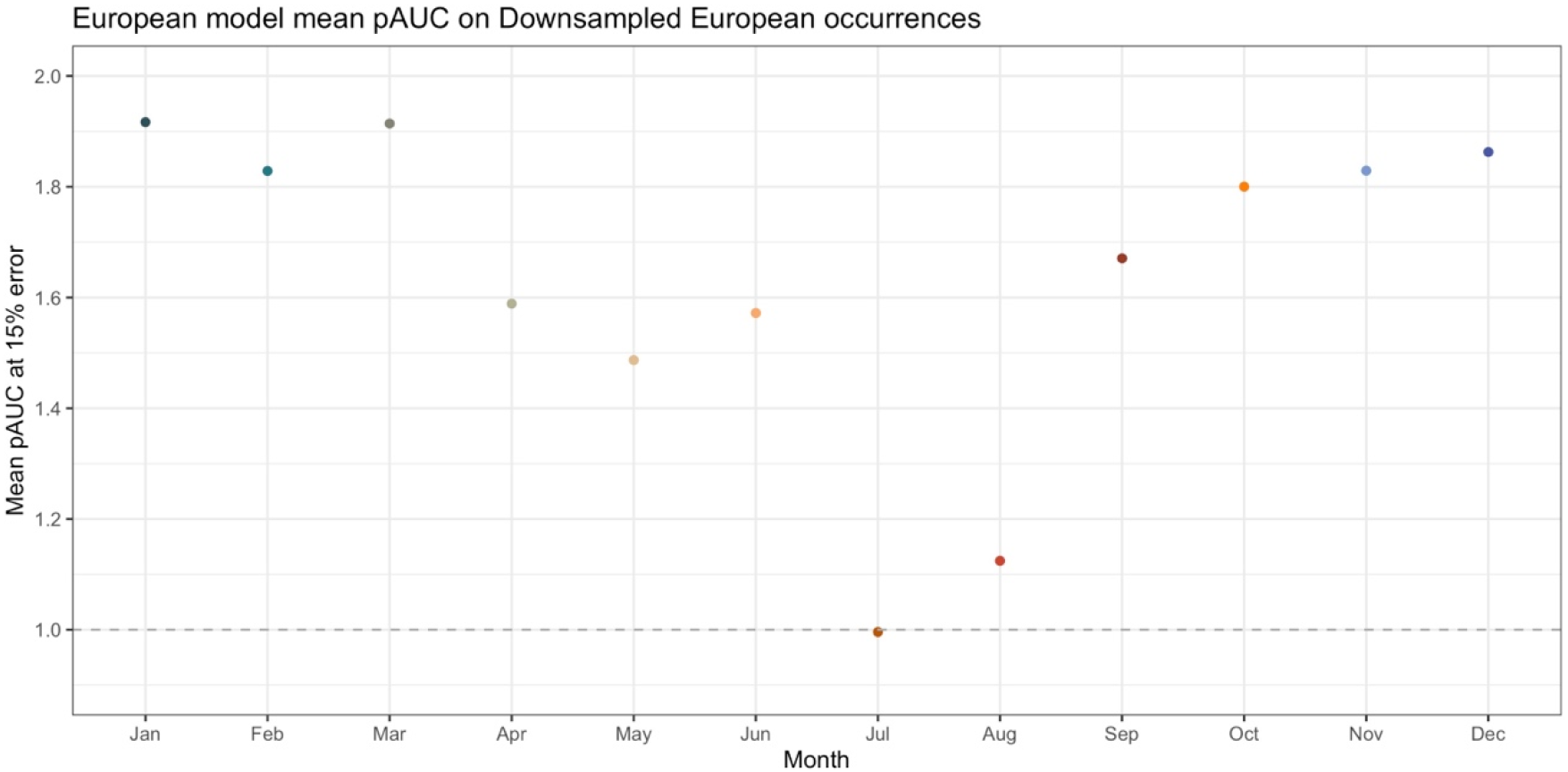
Predictive ability of European models with temporal downsampling on European occurrences. Each model was generated using 503 occurrences (the minimum number in the December models) per month.

**Supplementary Figure 11:**
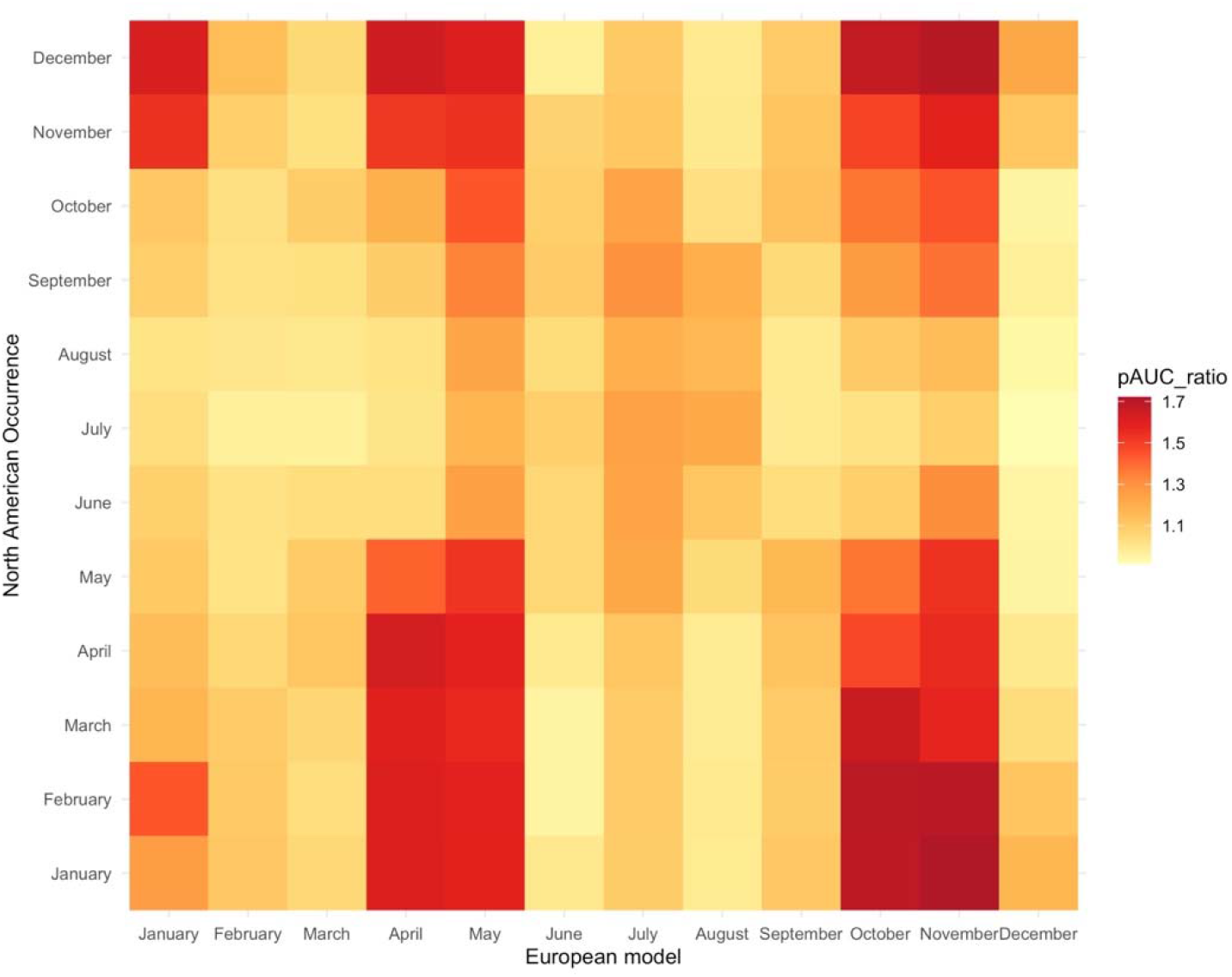
Predictive ability of European models with temporal down sampling on North American occurrences.

**Supplementary Figure 12:**
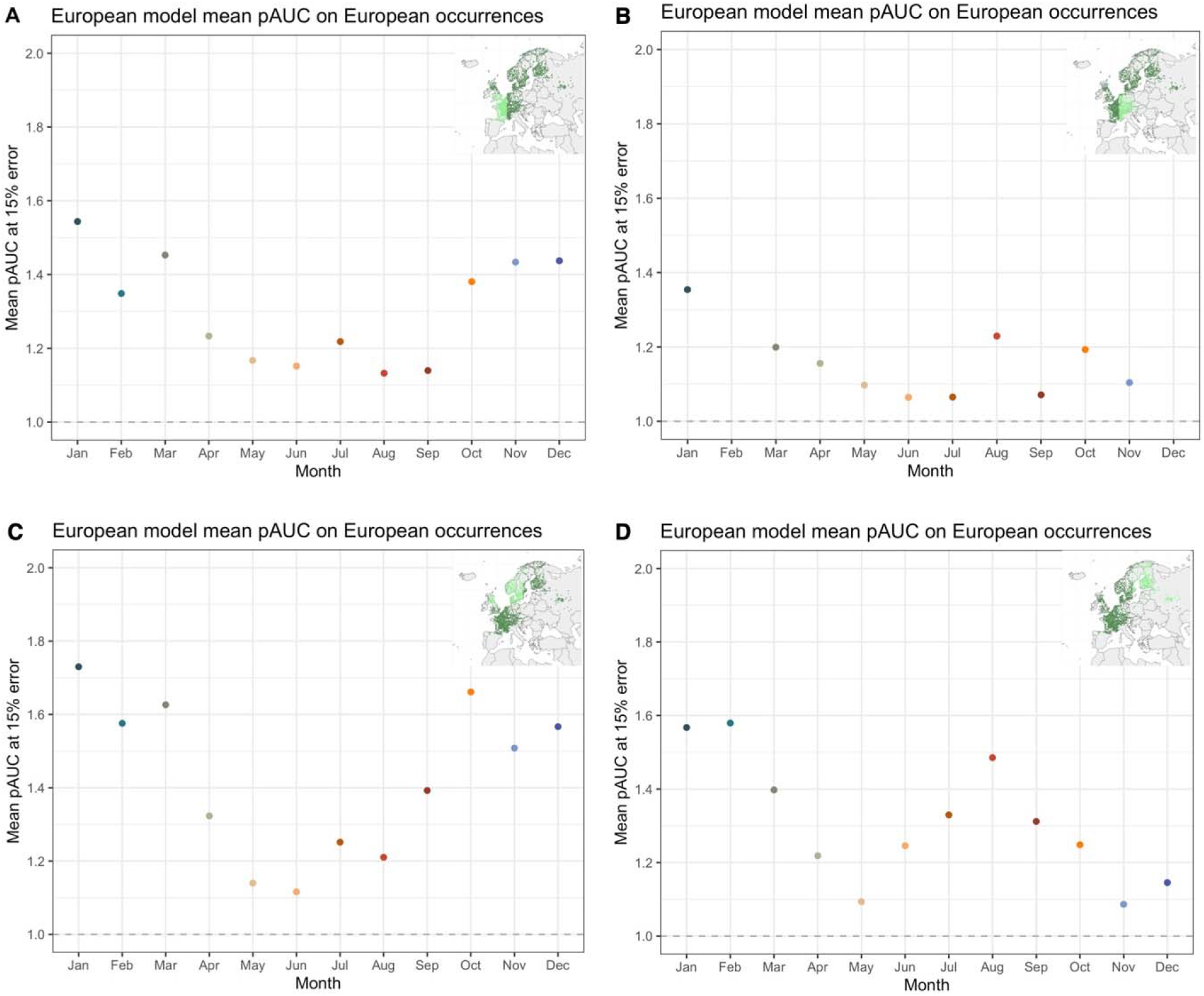
European model predictive ability on European testing points when all models have 25% data withheld for testing. Inset images illustrate the general geographic areas withheld for testing (using occurrences from August as an example) for each of the four bins.

